# Peak Anteroposterior Heel Slip Acceleration Predicts Lateral Trunk Flexion During Unexpected Slip Perturbations

**DOI:** 10.1101/2025.07.28.667233

**Authors:** Jonathan Lee-Confer, Dorie Chen, Karen Troy

## Abstract

Slip-induced falls are a major contributor to hip fractures and injury, particularly during sideways falls where lateral trunk flexion drives impact to the hip. While past studies have focused on sagittal-plane slip mechanics, the mechanisms of inducing frontal-plane trunk excursion remain poorly understood. This study investigated which slip foot kinematic variables predict lateral trunk flexion during unexpected slips. Twenty-six healthy young adults experienced an unexpected slip while walking in a laboratory setting. Peak anteroposterior (AP) heel slip distance, velocity, and acceleration, as well as mediolateral (ML) slip distance, velocity, and acceleration, were measured using three-dimensional motion capture. A stepwise multiple linear regression identified peak AP heel slip acceleration as the sole significant predictor of lateral trunk flexion (p = .026, R^2^ = 0.19). One-way repeated measures ANOVA and one-tailed paired t-tests further revealed that peak heel acceleration occurred significantly earlier than both peak heel velocity (p = .012) and the onset of lateral trunk flexion (p < .001), supporting its role as an early destabilizing mechanism. In contrast, neither velocity nor slip distance predicted lateral trunk flexion magnitude. These findings suggest that peak AP heel acceleration is associated with inducing lateral trunk flexion movements that likely precede lateral falls. Reducing heel acceleration through slip-resistant flooring or footwear, or enhancing compensatory strategies such as reactive arm movements, may reduce the likelihood of losing sideways balance. This work highlights the need for balance training interventions that emphasize rapid responses to early-phase slip dynamics.

**Highlights:**

- Every 1000 cm/s^2^ increase in AP heel acceleration leads to 6 degrees more lateral trunk flexion
- Heel acceleration occurs significantly earlier than heel velocity or the onset of lateral trunk flexion
- Findings highlight the importance in reducing forward heel acceleration to reduce sideways loss of balance

## Introduction

Slip-related falls are a major contributor to global injury rates and healthcare expenditures [1]. These incidents commonly result in injuries such as patellar and wrist fractures, as well as soft tissue damage to the shoulder. However, the most severe consequences are often hip fractures [2,3], which are associated with reductions in quality of life and require prolonged rehabilitation. Hip fractures are typically sustained when an individual falls sideways [4–6], impacting the greater trochanter of the femur. Despite the clinical significance, the biomechanical mechanisms underlying a sideways loss of balance during a slip remain poorly understood. Identifying these mechanisms is important to developing strategies aimed at preventing sideways falls and mitigating the risk of hip fractures.

The overall trajectory of the body during a slip has been a central focus in fall-related biomechanics research. Historically, the majority of studies have emphasized sagittal plane mechanics, largely due to early definitions of slip incidents that characterized them as backward losses of balance [7–13]. Much of the literature has investigated sagittal plane variables such as trunk kinematics [14], backward loss of stability [7,13,15], and angular momentum [16,17]. However, emerging research has begun to highlight the importance of frontal plane dynamics in slip-related falls [18–24]. For instance, one study reported that whole-body angular momentum in the frontal plane was approximately three times greater than that observed in the sagittal plane during slip events [24]. Additionally, several recent investigations have examined lateral trunk flexion and described the tendency of the body to fall laterally during a slip [18,23,25,26]. These findings suggest that frontal plane mechanics may play a more critical role in slip outcomes than previously recognized.

To date, the mechanisms underlying why the body rotates posteriorly or laterally during a slip remain poorly understood. Previous research has identified slip distance, or the displacement of the slipping foot, as a predictor of whether young adults experience a fall or successfully recover during a slip event [27]. Supporting this notion, Allin et al. (2018) demonstrated that specific features of foot kinematics such as peak anteroposterior (AP) and mediolateral (ML) heel slip speeds and slip distances accurately classified four distinct slip outcomes: recovery, feet-split falls, feet-forward falls, and lateral falls [23]. These findings highlight the potential of foot kinematics as a determinant of whole-body dynamics during slip events. Additionally, the earliest deviations in gait during a slip have been observed as reductions in vertical ground reaction force [28]. It may be that the faster the ground reaction force decreases during slip initiation, the faster anteroposterior heel acceleration may be.

What remains unknown is whether early-phase mechanical variables, such as heel acceleration, can better predict the onset of lateral trunk responses. Acceleration reflects the severity and abruptness of the perturbation at the foot-floor interface and may serve as a biomechanical contributor that initiates lateral trunk movements before foot displacement or velocity is evident.

To date, there is no study investigating the effects of peak anteroposterior heel acceleration on the mechanics of body rotations during a slip incident. Therefore, the purpose of this study was to investigate what foot kinematic variables during a slip would be a predictor of individuals exhibiting lateral trunk flexion from a slip incident. We hypothesized that peak anteroposterior heel acceleration of the slipped foot would best predict whether someone exhibits lateral trunk flexion during a slip incident.

## Methods

### Participants

26 healthy and young participants (Age: 25.46 ± 5.12 years old, Height: 170.42 ± 10.73 cm, Weight: 69.43 ± 15.63 kg, 14 Females and 12 Males, Table 1) were recruited for this IRB approved study. All participants signed a written informed consent after being provided with the scope of this study. Each participant was screened by an attending physician, and participants were excluded if they reported any musculoskeletal, cardiovascular or neurological conditions.

**Table 1.** Participant demographic characteristics (n = 26). Values for continuous variables are presented as mean ± standard deviation. Categorical variables are shown as counts and percentages.

| Variable | Mean ± SD |
| --- | --- |
| Age (years) | 25.46 ± 5.12 |
| Height (cm) | 170.42 ± 10.74 |
| Weight (kg) | 69.43 ± 15.63 |
| Sex (Male/Female) | 12 (48%) / 13 (52%) |

### Instrumentation

Detailed experimental methodology of this work has been previously described. All walking trials were performed on a designated walkway within the laboratory at the University of Illinois Chicago. A 1.22 m x 2.44 m x 0.63 cm Plexiglas sheet was embedded into the walkway. All participants were fitted into a full-body safety harness that supported the entire body weight of the participant (if needed) and prevented contact of the participants’ hands, knees, or buttocks with the floor should a fall occur. Participants were permitted and encouraged to partake in several practice walking trials across the walkway to acclimate to the harness. 3-D motion analysis was collected with eight motion capture camera system at 60 Hz (Motion Analysis, Santa Rosa, CA). Reflective joint markers were adhered to anatomical locations per the manufacturer’s guidelines to produce a 13-segment rigid body model to generate full-body joint kinematics (OrthoTrak, Motion Analysis, Santa Rosa, CA).

### Procedures

Participants were aware that they may be slipped on any walking trial, however participants were not aware of how many non-slip (control) walking trials would be conducted, nor which trial the slip would initiate. Furthermore, all participants were unaware of how a slip would be induced on the walkway. All participants were subjected to an unexpected slip perturbation by stepping onto a thin applied layer of mineral oil onto the Plexiglas when the participants were not looking.

### Data Analysis

Heel strike was defined as the moment when the Z-position of the heel marker was at its lowest point after the swing phase of gait. The X-position of the heel marker at heel strike was used as the initial starting point for the AP slip calculations and the Y-position of the heel marker at heel strike was used as the initial starting point for the ML slip calculations. AP slip distance was calculated by taking the displacement between the initial X-position and the final X-position after the foot stopped moving anterior from the slip. ML slip distance was calculated by taking the displacement between the initial Y-position and the final Y-position after the foot stopped moving laterally from the slip. For both directions AP and ML, heel slip velocity was calculated by the central difference method of position, and heel slip acceleration was calculated by the central difference method of velocity. Peak heel slip velocity and peak heel slip acceleration were the maximum values extracted from the beginning to the end of the slip for each of the respective variables. Peak lateral trunk flexion was defined as the angular excursion from initial trunk position at heel strike to the maximum lateral trunk angle during the slip. The onset timings for peak anteroposterior heel velocity and peak anteroposterior heel acceleration were defined as the time points after slip initiation when each variable reached its maximum value, while the onset of lateral trunk flexion was defined as the time point after slip initiation when the trunk angle first exhibited a clear directional change in the frontal plane. (Fig. 1).

**Figure 1.**
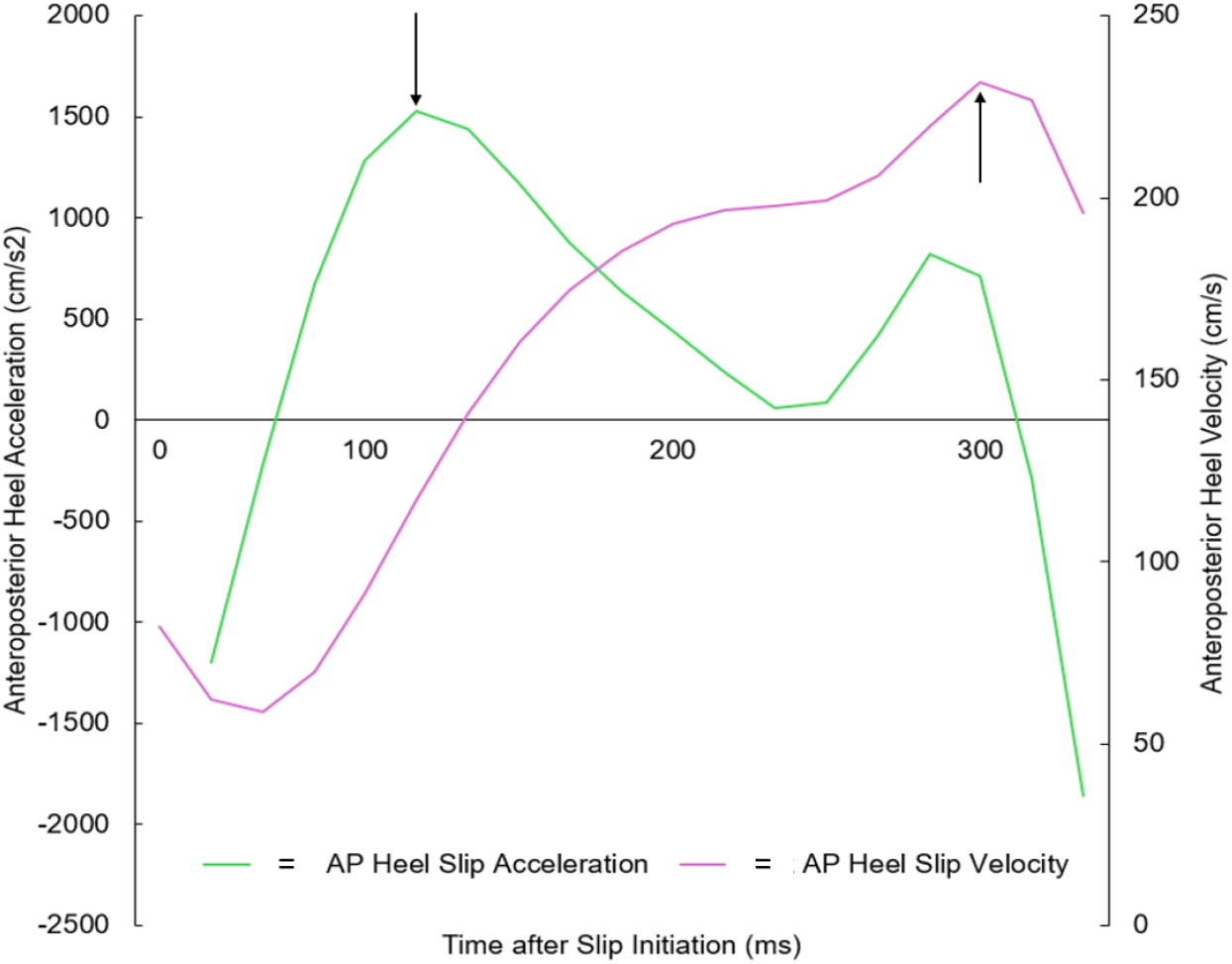
Representative graph illustrating an individual’s anterior-posterior (AP) heel acceleration (light green) and heel velocity (purple). Black arrows mark the respective peak values used to identify peak AP heel acceleration and peak AP heel velocity.

### Statistical Analysis

A stepwise multiple linear regression was selected for this exploratory analysis to identify predictor(s) of peak lateral trunk flexion from a larger set of foot kinematic variables during a slip. This approach allows for the systematic inclusion of predictors based on their statistical contribution to the model, reducing the risk of multicollinearity and model overfitting when theoretical justifications for variable inclusion are limited. Given the intercorrelations expected among slip distance, velocity, and acceleration metrics in both the AP and ML directions, a stepwise regression provided an objective method to isolate the strongest statistical contributor while controlling for redundancy among predictors. All variables were entered as candidates, and inclusion was based on p-value thresholds (entry: p < 0.05; removal: p > 0.10). To identify exploratory predictors of lateral trunk flexion, we conducted a stepwise multiple linear regression with peak lateral trunk flexion angle as the dependent variable and six slip-related kinematic variables of the foot (AP and ML slip distance, velocity, and acceleration) as candidate predictors. A one-way repeated measures ANOVA was conducted to assess the temporal sequence between heel kinematic events and trunk responses by comparing the timing of peak AP heel acceleration, peak AP heel velocity, and onset of lateral trunk flexion. One-tailed paired-samples t-tests were conducted to test the directional hypotheses of heel acceleration occurring earlier than heel velocity, and that both peak AP heel acceleration and peak AP heel velocity precede the onset of lateral trunk flexion. All assumptions were verified before analysis, and significance was set at α = 0.05. Analyses were performed using SPSS version 29 (IBM Corp., Armonk, NY)

## Results

The regression model identified peak anteroposterior (AP) heel slip acceleration as the sole significant predictor of peak lateral trunk flexion. The final model was statistically significant, F (1,26) = 5.60, p = .026, and explained approximately 18.9% of the variance in peak lateral trunk flexion (R^2^ = .189, adjusted R^2^ = .155).

Peak AP heel slip acceleration demonstrated a moderate and positive relationship with lateral trunk flexion (B = 0.006, SE = 0.002, standardized β = 0.435, t = 2.366, p = .026, Table 2). All other candidate predictors (AP heel slip distance, AP heel slip velocity, ML heel slip distance, ML heel slip velocity, and ML heel slip acceleration) were excluded from the model due to lack of statistical significance (p > .10). The range of p-values among the other candidate predictors were 0.301 to 0.924.

**Table 2.** Results from a stepwise multiple linear regression model identifying peak anteroposterior (AP) heel slip acceleration as the sole significant predictor of lateral trunk flexion during a slip. Coefficients (B), standard error (SE), standardized beta (β), t-statistic, and p-value are reported for the included predictor. All other candidate variables were excluded from the model due to lack of statistical significance (p > .10). The range of p-values among the other candidate predictors were 0.301 to 0.924.

| Predictor | B | SE(B) | B (Standardized) | t | p-value |
| --- | --- | --- | --- | --- | --- |
| AP Heel Slip Acceleration | 0.006 | 0.002 | 0.435 | 2.366 | .026 |
| Excluded Variables |  |  |  |  |  |
| AP Heel Slip Distance | - | - | - | - | > .10 |
| AP Heel Slip Velocity | - | - | - | - | > .10 |
| ML Heel Slip Distance | - | - | - | - | > .10 |
| ML Heel Slip Velocity | - | - | - | - | > .10 |
| ML Heel Slip Acceleration | - | - | - | - | > .10 |

Peak AP heel acceleration was the strongest correlate of peak lateral trunk flexion (r = 0.43, p = .026; see Fig. 1 and Fig. 2). Furthermore, when peak AP heel acceleration was stratified into quartiles, individuals in the lowest acceleration quartile exhibited the smallest lateral trunk flexion responses, while those in the highest quartile exhibited the greatest flexion (Fig. 3). This trend supports the positive linear association between heel acceleration and frontal-plane trunk motion observed in the regression analysis.

**Figure 2.**
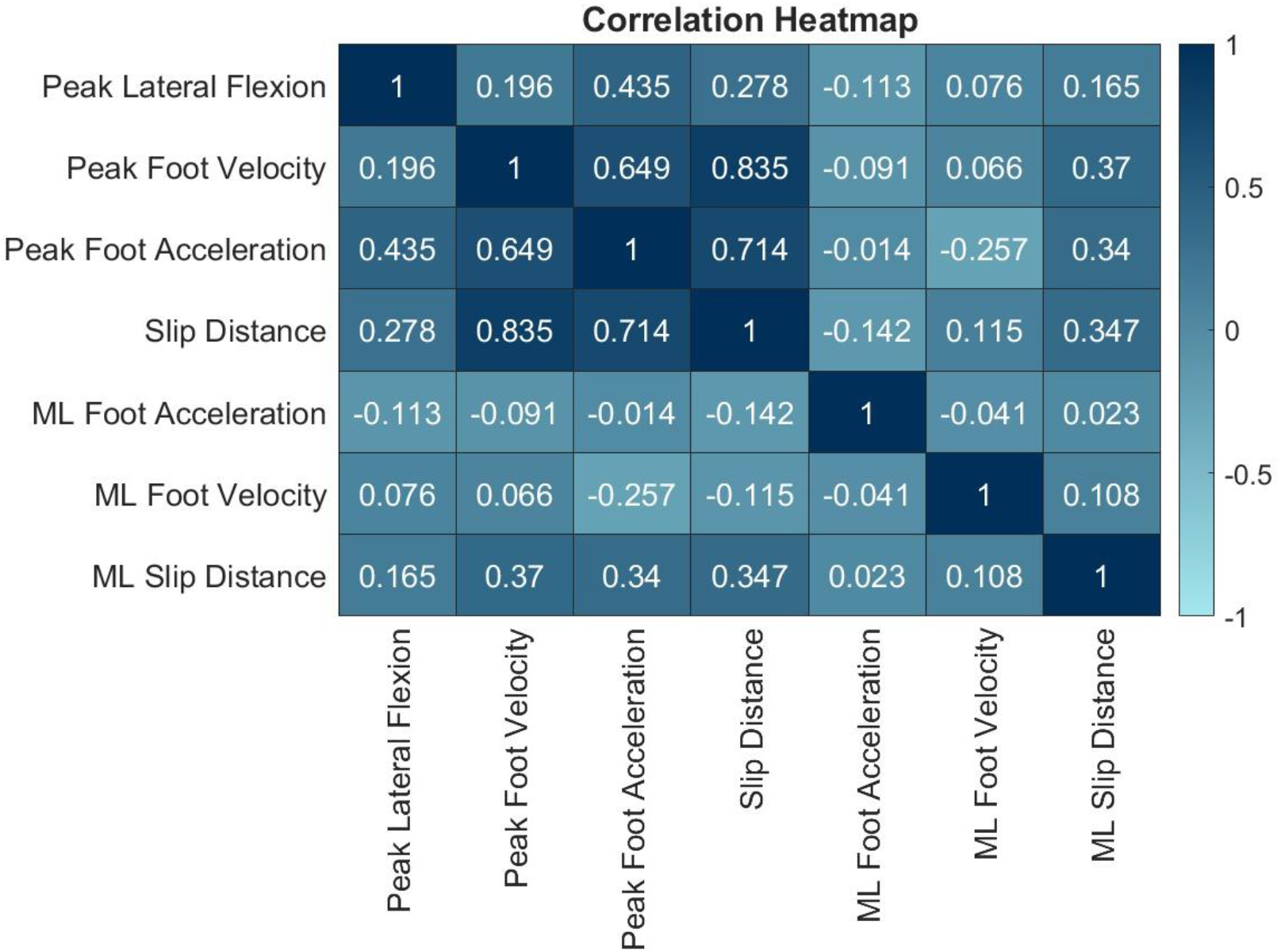
Pearson correlation coefficients among all foot kinematic variables and lateral trunk flexion. The heatmap displays pairwise correlation coefficients for six slip-related kinematic variables, peak AP and ML heel slip distance, velocity, and acceleration, as well as peak lateral trunk flexion. Darker colors represent positive correlations, while brighter colors represent negative correlations. AP heel slip acceleration shows the strongest positive correlation with lateral trunk flexion, consistent with regression results. Other variables show minimal or no linear relationship with trunk behavior.

**Figure 3.**
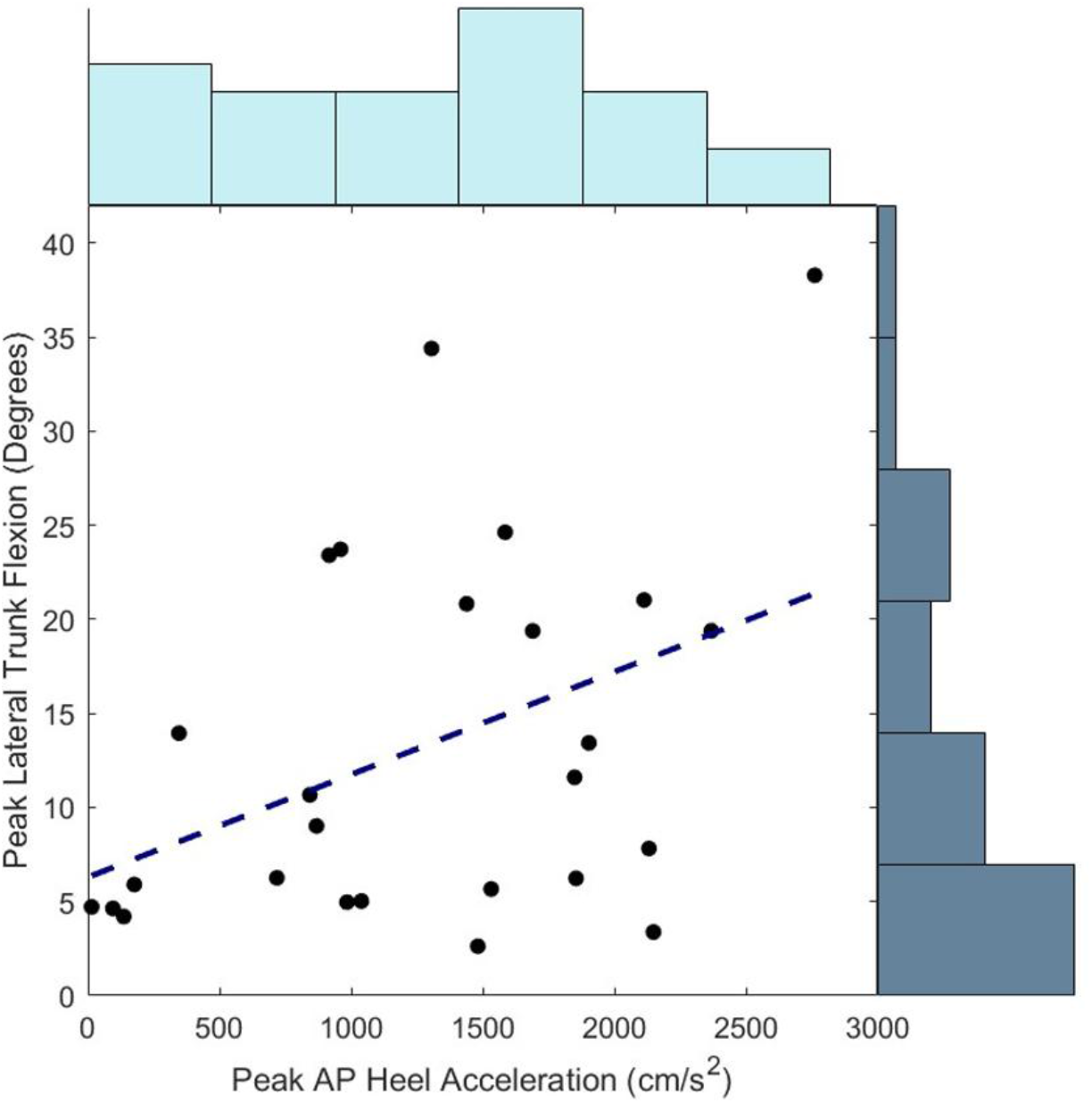
Relationship between peak anteroposterior (AP) heel slip acceleration and lateral trunk flexion during a slip. Each point represents data from a single participant. The bottom x-axis reflects peak AP heel slip acceleration (in cm/s^2^), and the bottom y-axis represents the corresponding peak lateral trunk flexion angle (in degrees). A positive linear relationship was observed, with the regression line defined by the equation y = 0.0057x + 5.60 (R^2^ = 0.189, p = .026). The distribution of the data for peak AP heel acceleration is on the top x-axis and the distribution of the data for lateral trunk flexion is on the right y-axis.

Collinearity statistics indicated no concerns, with a tolerance of 1.000 and a variance inflation factor (VIF) of 1.000 for the included variable. Residual diagnostics showed acceptable collinearity and variance proportions, with a condition index of 1.000.

A one-way repeated measures ANOVA was conducted to compare the timing of peak anteroposterior (AP) heel velocity, peak AP heel acceleration, and the onset of lateral trunk flexion following slip initiation. Multivariate tests indicated a significant main effect of event type on timing, Pillai’s trace = 0.500, F(2, 24) = 12.010, p < 0.001, partial η^2^ = 0.500; similar results were found using Wilks’ lambda. Mauchly’s test indicated that the assumption of sphericity was violated (W = 0.411, χ^2^(2) = 21.328, p < 0.001, and therefore a Greenhouse-Geisser correction was applied (ε = 0.629). The mean time to peak heel velocity was 286 ± 350 ms, the mean time to peak heel acceleration was 126 ± 85 ms, and the mean time to the onset of lateral trunk flexion was 256 ± 124 ms.

One-tailed paired-samples t-tests revealed that the time to peak AP heel acceleration (mean difference = -159.82 ms, SD = 339.11 ms) occurred significantly earlier than the time to peak AP heel velocity, t(25) = 2.403, p = 0.012. No significant difference was found between the time to peak heel velocity and the onset of lateral trunk flexion (mean difference = 29.69 ms, SD = 395.66 ms), t(25) = 0.383, p = 0.353. However, the time to peak heel acceleration occurred significantly earlier than the onset of lateral trunk flexion (mean difference = –130.13 ms, SD = 164.39 ms), t(25) = –4.036, p < 0.001.

## Discussion

The purpose of the current study was to identify a slip severity metric that would predict lateral trunk flexion during a slip incident. Among the six candidate predictors of peak anteroposterior (AP) heel slip distance, velocity, and acceleration, and peak mediolateral (ML) heel slip distance, velocity, and acceleration, only peak AP heel slip acceleration was a statistically significant predictor of lateral trunk flexion. This finding supports the hypothesis that slip severity, particularly the acceleration of AP foot motion after slip onset, plays a major role in contributing to lateral trunk flexion.

Peak AP heel slip acceleration likely represents the earliest and most intense perturbation to postural stability following heel strike during a slip incident. As individuals transfer weight onto the slipping limb, a sudden acceleration of the foot in the anterior direction causes the supporting limb to slide out from under the body’s center of mass. This forward translation of the stance limb can result in a pelvic drop on the side of the slipped foot and cause compensatory lateral flexion of the trunk toward the slipping side. The one-tailed paired-samples t-tests showed that peak AP heel acceleration occurs significantly earlier than both peak AP heel velocity and the onset of lateral trunk flexion. This temporal sequence strengthens the argument that heel acceleration serves not only as a statistical predictor but also as the mechanical variable most associated with inducing lateral trunk flexion responses.

In contrast, peak AP heel slip velocity and slip distance were not significant predictors of lateral trunk flexion. While velocity is often used to classify fall types [23], it reflects a later stage of the slip response, after rapid destabilizing accelerations have already taken effect. Similarly, slip distance indicates the total movement but not the initial destabilizing impulse that initiates lateral trunk flexion. These findings indicate that early mechanical foot disturbances induce lateral trunk flexion excursions as AP heel acceleration is the only variable that precedes lateral trunk flexion.

The ML slip variables (distance, velocity, and acceleration) also failed to predict lateral trunk flexion, despite an intuitive connection between frontal plane foot motion and trunk lateral flexion. This may be due to the relatively small or inconsistent ML slip severities observed in this study. The slips in this dataset were largely dominated by AP movement, which is expected, limiting the mechanical influence of ML foot motion on lateral trunk behavior. Furthermore, other work has shown that hip abduction does not significantly differ between slip trials and regular walking trials [29], supporting the idea that there is less frontal plane motion of the legs during a slip incident, and therefore less ML foot kinematics. Lastly, ML movements directed laterally would create a ground reaction force that would push the body towards the midline, or more upright, and not contribute to further ipsilateral lateral movements of the trunk.

While our findings emphasize AP heel acceleration as the key predictor, prior research by Allin et al. (2018) identified ML foot kinematics, particularly ML velocity and distance, as significant factors distinguishing recovery from various fall outcomes, including lateral falls. The contrast in our findings may be explained by differences in study design as Allin et al. used group-level discriminant analyses to classify fall types [23], whereas our study modeled continuous lateral trunk flexion to directly assess how slip dynamics predict lateral trunk flexion responses. Additionally, our inclusion of ML acceleration, which Allin et al. did not assess, may offer a more temporally sensitive measure of slip onset dynamics. Together, these contrasts suggest that while ML displacement and velocity can help classify overall fall outcomes, peak AP heel acceleration may contribute to driving the lateral flexion trunk movements that precede full-body loss of balance.

The current findings suggest that reducing heel slip acceleration may be a promising approach to minimize the likelihood of sideways falls. Since heel acceleration is the earliest and strongest predictor of resultant lateral trunk flexion, moderating AP heel acceleration through improved footwear [30–33], slip-resistant flooring [34–36], improving detection [37,38] or response time to slips, or strengthening of the hamstrings [39] could reduce peak AP heel acceleration, thus reducing lateral trunk flexion, and ultimately reducing the risk of a lateral fall that could lead to a hip fracture. This study corroborates findings previously that reported hamstring fatigue resulted in an increased slip severity [40] due to the inability to pull the slipped foot backwards after its forward displacement from the slip.

Despite the novel findings, several limitations should be considered. The sample consisted of healthy young adults, limiting generalizability to older adults or clinical populations with different neuromuscular responses. Only the initial slip trial was analyzed for each participant, which may not capture adaptation effects or within-subject variability. This study does not have electromyography data of the hamstrings to determine if hamstring activation has the ability to reduce heel slip acceleration. Finally, the variance explained in this study was low, suggesting further studies needing to include more variables such as muscle response times, arm reactions, muscle strength and more.

Future studies should expand to older or balance-impaired populations, who are more vulnerable to hip fractures from lateral falls. Research should also explore multiplanar trunk dynamics, angular momentum, and muscle activation patterns (e.g., via EMG) to better understand the neuromechanical strategies that respond to high slip accelerations. These insights can inform the design of more responsive assistive devices, perturbation-based balance training programs, and wearable sensor systems aimed at detecting and mitigating slip-induced destabilization in real time.

## Ethics Statement

This study was conducted in accordance with the ethical standards of the Declaration of Helsinki and was approved by the Institutional Review Board at the University of Illinois Chicago. All participants provided written informed consent prior to participation after receiving a full explanation of the study procedures and potential risks. Participants were screened for eligibility by a licensed physician, and individuals with musculoskeletal, cardiovascular, or neurological conditions were excluded. Participant safety was ensured through the use of a full-body harness to prevent injury during slip trials.

## Conflict of Interest Statement

The authors declare no commercial or financial relationships that could be construed as a potential conflict of interest.

## Funding Sources Statement

This research received no specific grant from any funding agency in the public, commercial, or not-for-profit sectors.

## Author Contribution Statement

Jonathan S. Lee-Confer conceptualized the study, performed data analysis, interpreted the results, and drafted the manuscript. Dorie Chen assisted with data processing, figure generation, and manuscript editing. Karen Troy contributed to study design, curation of the data, interpretation of findings, and revision of the manuscript. All authors reviewed and approved the final version of the manuscript.

## Data Availability Statement

The datasets generated and analyzed during the current study are available from the corresponding author on reasonable request.

## Notes

### Competing Interest Statement

The authors have declared no competing interest.

